# Engineered probiotics target granulomas to treat mycobacterial infection

**DOI:** 10.64898/2026.10.27.762313

**Authors:** Yu-Yu Chen, Tetsuhiro Harimoto, Gaëlle Guiewi Makafe, Courtney Coker, Hannah Kim, Jongwon Im, Yuanqi Jing, Hayoung Jin, Xiaoyu Hu, Sharicka Zutshi, Mia Ross Raneri, Brian Weinrick, Tal Danino

## Abstract

Granulomas are a pathological niche for mycobacteria that remain difficult to access with conventional therapies. Here we show that probiotic *Escherichia coli* Nissle 1917 (EcN) infiltrates and colonizes granulomas and can be engineered as a delivery system that senses the local microenvironment and releases antimycobacterial payloads. Specifically, we leveraged the hypoxic microenvironment of necrotic granulomas to restrict bacterial growth, and developed a dual-mode lysis circuit combining intracellular and extracellular killing for sustained payload release with limited evolutionary escape. We further engineered EcN to express a mycobacteriophage-derived enzyme, generating a strain that suppressed drug-sensitive and drug-resistant mycobacteria in vitro and potentiated conventional antibiotic activity. Engineered probiotics also reduced mycobacterial burden in vivo, and in a murine model of rifampicin-resistant infection, combination treatment with pretomanid achieved greater bacterial reduction than either monotherapy. Together, these results establish a synthetic biology framework for targeting granulomatous infections with programmable living microbes.

## Background

Engineered microbial systems are emerging as living therapeutic modalities capable of sensing physiological environments and delivering payloads in vivo^1,2^. One key advantage is the natural capacity of bacteria to colonize native tissue niches — such as the gastrointestinal tract, lung, and skin — where they can be engineered to suppress pathogenic organisms^3–6^ or modulate host physiology^7–9^. Within these niches, synthetic biology–enabled gene circuits further provide precise environmental sensing, recording, computation, and regulated therapeutic output^10–14^. Bacteria have been shown to preferentially proliferate in environments such as hypoxic and necrotic tumor cores, where they can be engineered to detect local cues and modulate the microenvironment, with several strains advancing to clinical evaluation^15–20^.

Granulomas represent a similarly structured but unexplored niche for microbial therapies. These organized immune aggregates form during chronic infection to contain persistent pathogens and are defined by hypoxic, necrotic cores, restricted vascularization, and limited immune accessibility^21–24^. Such environments are central to a range of granulomatous diseases, including tuberculosis (TB), nontuberculous mycobacterial infections, and other chronic infections such as salmonellosis, listeriosis, and selected fungal or parasitic diseases. Among these, TB remains the most clinically significant, with *Mycobacterium tuberculosis (M. tuberculosis)* infecting approximately one-quarter of the global population and causing over a million deaths annually. Despite recent advances in TB therapeutics, including new agents for multidrug-resistant disease, treatment remains limited by prolonged systemic administration, toxicity, and the emergence of resistance. Alternative strategies are being developed—including bacteriophage therapy^25^, immunomodulatory biologics^26^, and cellular therapies such as mesenchymal stem cells^27^. Critically, these approaches do not overcome the fundamental challenge of poor drug penetration into granulomas, where structural and microenvironmental barriers limit therapeutic access and promote pathogen persistence. These limitations motivate the development of engineered bacteria as localized, programmable therapeutics for granulomatous infections.

Here, we engineered a probiotic bacteria, *E. coli* Nissle 1917 (EcN), to selectively infiltrate granulomas and locally produce and release antimycobacterial payloads. We designed hypoxia-responsive gene circuits and quorum-regulated lysis systems to create a strain that (1) preferentially proliferates under granuloma-associated hypoxia, (2) undergoes lysis for sustained therapeutic release, and (3) expresses antimicrobial proteins capable of suppressing both drug-sensitive and drug-resistant mycobacteria (**Fig. 1a**). Together, this work establishes a new strategy for mycobacterium therapy and provides a modular framework for targeting other granulomatous infections.

**Fig. 1.**
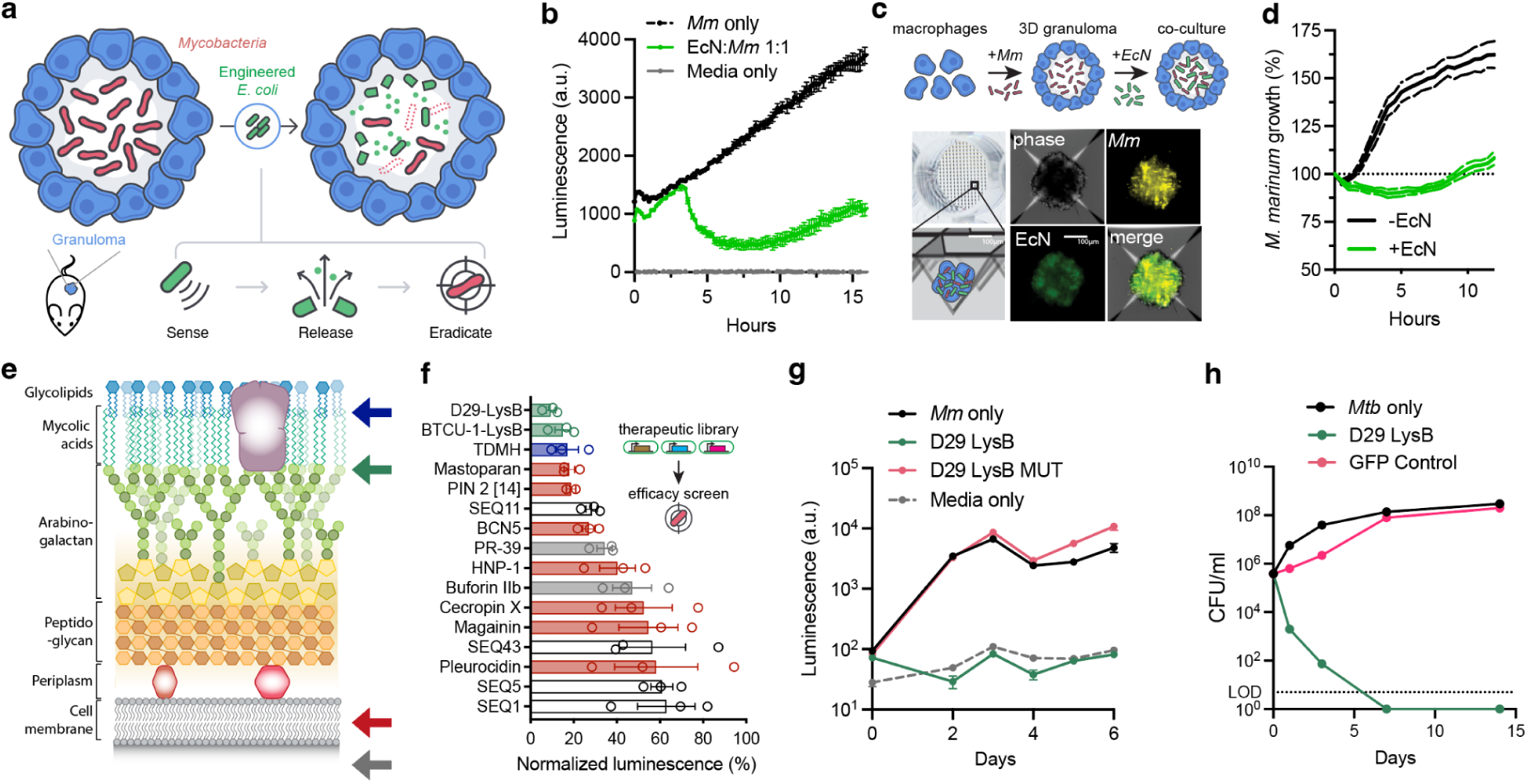
Engineered probiotic *E. coli* Nissle 1917 suppresses mycobacteria in granuloma models. **a**, Schematic of engineered *E. coli* Nissle 1917 (EcN) for granuloma therapy. EcN colonizes granulomas, senses hypoxia and releases payloads via a lysis circuit. **b,** Co-culture assay showing EcN suppresses luminescent *M. marinum* (*Mm*-lux) at 1:1 ratio. *Mm*-lux increased ∼8-fold over 15 hours in monoculture but declined within 3 hours of co-culture, consistent with interspecies competition. **c,** Schematic and representative images of 3D macrophage-based granuloma spheroids infected with *M. marinum*-tdTomato (*Mm-*tdT) and co-cultured with EcN expressing GFP for 6 hours. Microscopy on the following day showed EcN colocalization with infected granulomas and reduction of mycobacterial signal. Scale bar, 100 µm. **d,** Quantification of *M. marinum* growth dynamics in granuloma spheroid with (+EcN) or without (−EcN) co-culture. EcN transiently suppresses growth but does not sustain clearance beyond ∼5 hours, indicating that competition alone is insufficient (n = 6 spheroids per condition). **e,** Mycobacterial cell wall diagram highlighting AMP target sites. **f,** AMP library expressed in EcN screen against *Mm*-lux; bar color indicates mechanism of action target: mycolic acid (blue), arabinogalactan (green), cell membrane (red), intracellular machinery (grey), and unknown (white). **g,** *Mm*-lux luminescence over 6 days following treatment with D29 LysB or catalytic mutant (D29 LysB MUT) EcN lysate. D29 LysB sustained complete suppression, whereas the mutant had no effect, confirming enzyme-dependent inhibition. **h,** CFU/mL of *M. tuberculosis* cultures treated with D29 LysB- or GFP-EcN lysate. D29 LysB sterilized cultures within 7 days, whereas GFP control lysate had no effect. LOD, limit of detection. Data represent mean ± s.e.m. (b–g) or SD (h); n = 3 biological replicates unless noted.

## Results

To determine whether EcN colonization alone could suppress mycobacterial growth and provide a baseline for subsequent therapeutic engineering, we first evaluated competitive interactions between bacterial species. We used *Mycobacterium marinum* (*M. marinum*), a BSL-2 surrogate for *M. tuberculosis* that shares its virulence and granuloma-forming pathology while enabling rapid experimental evaluation^28^. EcN expressing GFP (EcN-GFP) was co-cultured with *M. marinum*-expressing a luminescent reporter (*Mm*-lux; *luxCDABE*), enabling real-time tracking of each population. EcN, with a doubling time of ∼20 minutes in LB broth, rapidly outcompeted *Mm*-lux, which divided roughly every 6 hours in 7H9 liquid culture. In monoculture, *Mm*-lux luminescence increased about 8-fold over 15 hours, but declined sharply within 3 hours when co-cultured with EcN at a 1:1 ratio, consistent with growth inhibition through interspecies competition (**Fig. 1b**). To evaluate these interactions in a more physiologically relevant context, we established a high-throughput 3D granuloma co-culture platform that forms multicellular spheroids composed of macrophages and pathogenic mycobacteria, enabling continuous co-culture with EcN and time-lapse imaging of all three populations within a defined microenvironment (**Fig. 1c**)^29,30^. In this 3D granuloma model, *M. marinum* growth was initially suppressed following EcN colonization but resumed expansion after ∼5 hours (**Fig. 1d**), indicating that competition alone provides transient suppression but is insufficient for sustained mycobacterial control.

To convert the transient suppression observed with EcN alone into sustained antimycobacterial activity, we engineered EcN to locally produce antimicrobial peptides and proteins (AMPs), which are otherwise limited by instability, poor bioavailability, and toxicity when administered systemically^31^. To identify potent payloads, we constructed a library with 16 diverse AMPs targeting key mycobacterial structures, including the cell envelope and intracellular machinery (**Fig. 1e**). Each AMP was expressed from a high-copy ColE1 plasmid to ensure robust production while maintaining tolerable metabolic burden (**Supplementary Table 1**). Screening of crude lysates against *Mm*-lux identified multiple payloads with strong, dose-dependent activity; D29 LysB, BTCU-1 LysB, and TDMH achieved the greatest growth suppression relative to EcN-GFP control (**Fig. 1f**). Among these, D29 Lysin B (D29 LysB)—an esterase from mycobacteriophage D29 that cleaves the linkage between mycolic acids and arabinogalactan^32^—showed the strongest activity^33^. Mutation of the catalytic serine to alanine (D29 LysB MUT S82A) abolished activity without affecting expression (**Fig. 1g**)^32^. Lysates from D29 LysB-expressing EcN suppressed *M. marinum* growth for up to six days, whereas the mutant had no effect, confirming enzyme-dependent inhibition. A C-terminal His-tag enabled detection of both variants at the expected molecular weight (28.5 kDa; **Supplementary Fig. 1**). Finally, we tested whether this activity extended to *M. tuberculosis* (*M. tb*) in vitro, where EcN alone showed minimal inhibition, whereas D29 LysB-expressing EcN rapidly suppressed *Mtb* and sterilized cultures within 7 days (**Fig. 1h**).

### A dual-lysis circuit sustains payload release

Efficient therapeutic delivery requires controlled release of payloads from colonizing bacteria. A previously reported synchronized lysis circuit (SLC) could trigger payload release^17^, but incomplete lysis and evolutionary escape limited release efficiency and consistency. To address these limitations, we engineered a dual-mode lysis circuit (DUEL; **<u>du</u>**al **<u>E l</u>**ysis) that couples a previously described intracellular mechanism with a new extracellular killing mechanism to amplify payload release and suppress escape variants (**Fig. 2a**, **Supplementary Fig. 2**). DUEL combines quorum-controlled expression of the phage lysis gene ɸX174E (E) to trigger primary lysis, which releases colicin E1 (CE1) to induce secondary lysis in neighboring cells to propagate lysis across the bacterial population. To tune lysis dynamics, we constructed DUEL strains expressing CE1 under constitutive promoters of increasing strength on a medium-copy p15a plasmid (DUEL117, DUEL114 and DUEL119; **Supplementary Table 2**). These DUEL strains exhibited repeated lysis cycles and limited population overgrowth, whereas a control without CE1 (SLC) underwent a single lysis event followed by rapid expansion due to evolutionary escape. Increasing CE1 expression enhanced growth suppression, with DUEL117 maintaining growth prior to lysis, DUEL114 exhibiting stronger lysis, and the highest-expression variant (DUEL119) being nonviable (**Fig. 2a**). Time-lapse microscopy of DUEL117 revealed stepwise release of sfGFP corresponding to successive lysis events (**Fig. 2b, Supplementary Movie 1**), and extracellular GFP accumulation was significantly increased relative to non-lytic and SLC controls (**Fig. 2c**). To improve genetic stability, we integrated the DUEL117 circuit into the EcN genome at two chromosomal loci: *cspA* and *ompW*^34,35^. The resulting strain (i-DUEL) maintained stable, repeated lysis and sustained payload release (**Fig. 2d**), establishing a genetically stable platform for subsequent therapeutic studies.

**Fig. 2.**
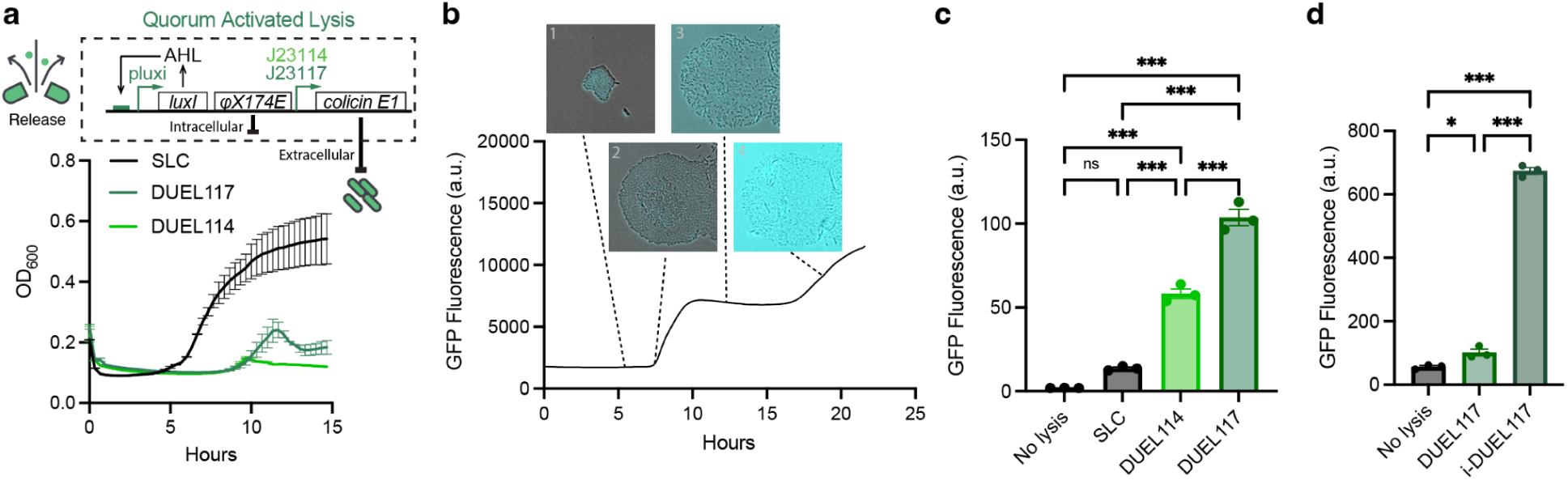
Quorum-activated DUEL-lysis circuit enables multiple and sustained release of payloads. **a**, The dual E lysis (DUEL) circuit couples luxI/luxR quorum-sensing to expression of the bacteriophage lysis gene (ϕX174E) and the extracellular lysis colicin E1. Growth dynamics (OD_600_) of strains carrying the synchronized lysis circuit (SLC; black) or DUEL constructs with varying CE1 expression levels (J23117 and J23114 promoters; dark green and light green respectively) demonstrate multi-cycle lysis and growth regulation while SLC undergoes a single lysis event followed by evolutionary escape and regrowth. **b,** Representative phase and fluorescence images taken by time-lapse microscopy showing sequential bacterial lysis in DUEL117 strain, corresponding to stepwise release of the reporter GFP diffusing into the medium. **c,** Quantification of extracellular GFP fluorescence scales across CE1 expression level circuit designs. **d,** Comparison of total GFP release between plasmid-based and double genomically integrated DUEL117 circuit (i-DUEL), demonstrating stable and efficient release from the integrated strain. Data represent mean ± s.e.m.; n = 3 biological replicates (a, c, d); one-way ANOVA with Tukey’s multiple comparisons test (c, d); \**P* < 0.033, \*\**P <* 0.002, \*\*\**P <* 0.001; ns, not significant.

### EcN infiltrates and colonizes granulomas in vivo

To determine whether EcN strains can infiltrate granulomatous lesions in vivo, we established local and systemic *M. marinum* (*Mm-*FLuc; firefly luciferase) infection models in C3HeB/FeJ mice. Intravenous infection generated cutaneous granulomas in the tails and footpads within two weeks, which was confirmed by *Mm*-FLuc signal from within necrotic regions (**Supplementary Fig. 3**). Localized footpad injection produced granulomas within one week, which was more tolerable and enabled faster experimental iteration (**Supplementary Fig. 4)**. Next, we intravenously administered EcN-lux after 14 days to mice infected intravenously with *Mm*-FLuc, and tracked both populations using their orthogonal bioluminescent reporters. EcN signals localized to granuloma-bearing tail regions and colocalized with *Mm* (**Fig. 3a**). Immunofluorescence of harvested tissues three days after injection confirmed EcN within caseous granulomas containing *Mm* (**Fig. 3b, Supplementary Fig. 5**), demonstrating its ability to infiltrate mycobacterium-infected lesions in vivo. Immunohistochemistry staining of direct injection into the mice tail with either *Mm* or EcN demonstrated that the specific antibodies can stain for either bacterial species with minimal cross-reactivity (**Supplementary Fig. 6**). The luminescent EcN signal in the tail granulomas continued to increase over the next three days, and high concentrations of viable EcN (∼10^7^ CFU/g) were recovered from granulomatous tissues at day three (**Fig. 3c**). Consistent colonization was also observed in the localized footpad infection model of *Mm*-FLuc, where systemically administered EcN-lux similarly accumulated within granulomas after three days (**Supplementary Fig. 7**). To our knowledge, these findings provide the first evidence that non-pathogenic bacteria EcN colonizes granulomatous tissues in vivo.

**Fig. 3.**
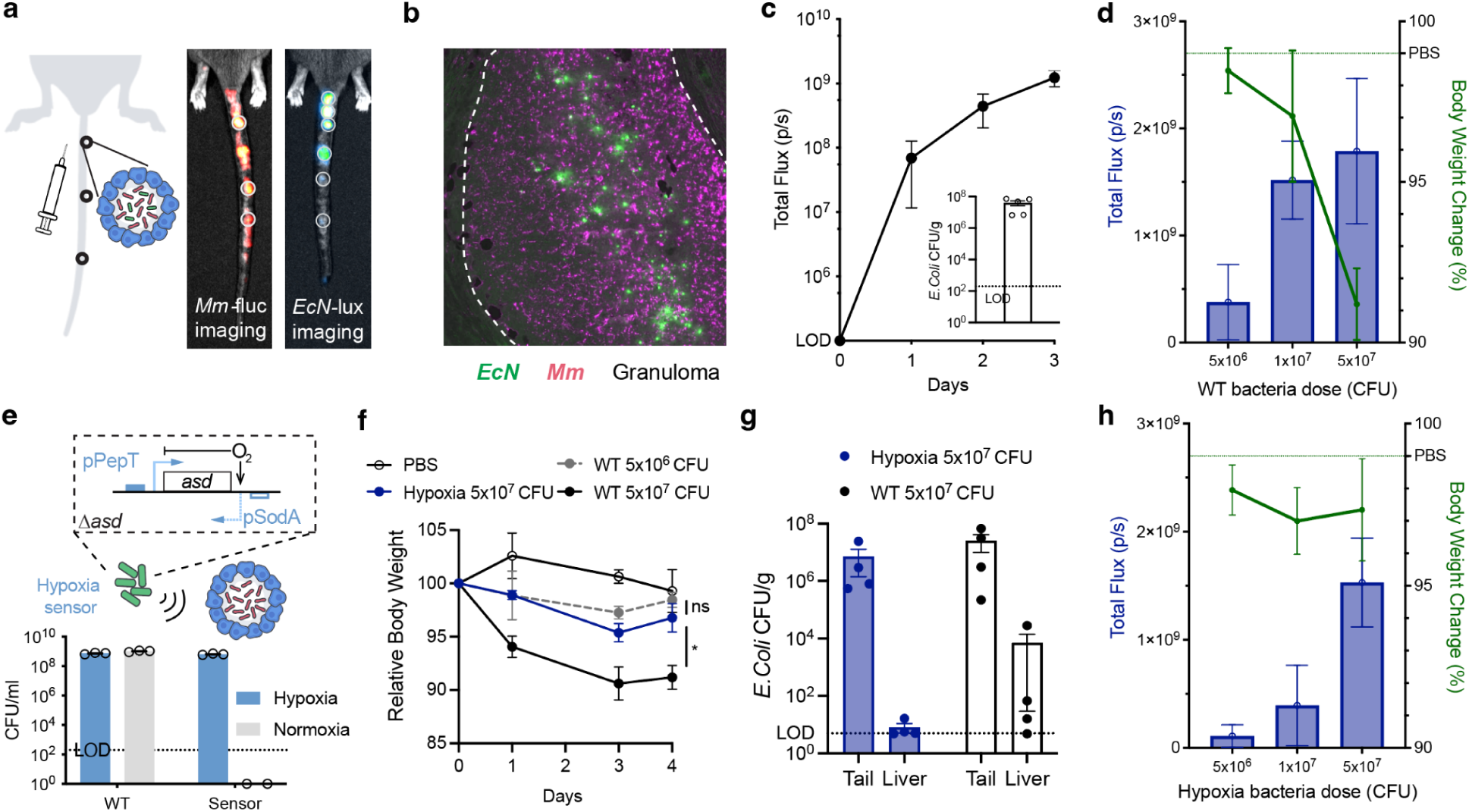
Hypoxia-sensing EcN selectively colonizes granulomas and reduces off-target accumulation in vivo. **a**, IVIS imaging of C3HeB/FeJ mice infected intravenously with *M. marinum* (*Mm*-FLuc) showing EcN-lux colocalization with tail granulomas 3 days post-injection. **b,** Multiplexed immunofluorescence showed EcN (anti-LPS) colocalizes within necrotic granulomas containing *M. marinum* (anti-mycobacterium) 3 days after systemic EcN administration. **c,** Longitudinal EcN-lux luminescence over 3 days shows progressive colonization; inset, viable EcN recovered (CFU/g) in tail tissue at day 3. **d,** Effect of varying EcN dose on luminescence intensity (left axis; blue) by measuring total flux [photons per second (p/s)] and percent body weight change (right axis; green) by day 4. **e,** Schematic of the hypoxia biosensor strain (hEcN) containing an *asd* knockout driven by the hypoxia-inducible pPepT promoter. Bacteria CFU/mL count showed selective hEcN growth under hypoxia vs normoxia. **f,** Relative body weight over 4 days following intravenous administration of PBS, WT EcN-lux (5×10^6^ and 5×10^7^ CFU), or hEcN-lux (5×10^7^ CFU). hEcN-treated mice maintained body weight comparable to PBS, whereas WT EcN induced significant weight loss. **g,** Biodistribution of WT EcN and hEcN in tail granulomas and liver on day 4. hEcN colonized granulomas at levels comparable to WT EcN but was undetectable in the liver. **h,** Effect of varying hEcN dose on luminescence intensity and body weight change by day 4. Higher hEcN doses increased granuloma colonization with better tolerability compared to equivalent WT doses. Data represent mean ± s.e.m. (c, d, f, g, h); n = 5 mice per group (c), n = 4 mice per group (d, f, g, h), n = 3 biological replicates (e). Statistical analysis was performed using two-way ANOVA (f); \**P* < 0.033; ns, not significant.

We evaluated EcN colonization across delivery routes and host backgrounds. With intravenous delivery of EcN, colonization was dose-dependent, with 5×10^6^ CFU representing a threshold for detectable localization. While a higher dosage at 5×10^7^ CFU enhanced EcN colonization, mice experienced increasing toxicity, indicated by body weight loss (**Fig. 3d)**. To exclude the possibility that tail granuloma colonization resulted from local exposure during tail-vein administration, we also delivered EcN through the retro-orbital (RO) route. EcN similarly localized to granulomas following RO administration in both C3HeB/FeJ and C57BL/6J mice, demonstrating colonization across delivery routes and host backgrounds (**Supplementary Fig. 8**). To improve tolerability, we engineered a hypoxia-dependent EcN strain (hEcN) by placing the essential gene *asd* under control of a hypoxia-responsive promoter^36^. hEcN retained robust colonization under hypoxic conditions in the 3D-granuloma model, confirming selective growth in low-oxygen environments (**Fig. 3e**). Consistent with this, hEcN administration in vivo maintained granuloma colonization while minimizing weight loss due to bacterial burden (**Supplementary Fig. 9**). Following intravenous administration in C3HeB/FeJ mice, hEcN exhibited improved tolerability, shown by the reduced weight loss (**Fig. 3f**), and decreased off-target accumulation in the liver compared to EcN (**Fig. 3g**), along with an improved dose–toxicity relationship (**Fig. 3h**). Together, these results demonstrate that hypoxia-dependent growth preferentially restricts EcN to granulomatous environments in vivo, resulting in an enhanced therapeutic index.

### hit-DUEL suppresses mycobacterial growth in vitro

We combined all three modules: <u>h</u>ypoxia sensing, <u>i-</u>DUEL lysis, and <u>t</u>herapeutic expression (D29 LysB), into a single strain, <u>hit</u>-DUEL EcN (**Fig. 4a**). To assess therapeutic efficacy, we treated luminescent *M. marinum* (*Mm*-lux; *luxCDABE*) with increasing concentrations of hit-DUEL lysate. hit-DUEL lysate suppressed *Mm*-lux growth in a dose-dependent manner, achieving complete killing at high concentrations; however, at concentrations more likely achievable within granulomas, growth suppression was only transient. Similarly, pretomanid (PA-824), a clinically used antibiotic for multidrug-resistant TB, achieved only temporary inhibition as a monotherapy at subtherapeutic concentrations^37–39^. We therefore tested whether combining hit-DUEL lysate with subtherapeutic PA-824 could achieve sustained growth control. Combination treatment led to sustained suppression of *Mm*-lux growth (**Fig. 4b**), demonstrating potentiation of antibiotic activity.

**Fig. 4.**
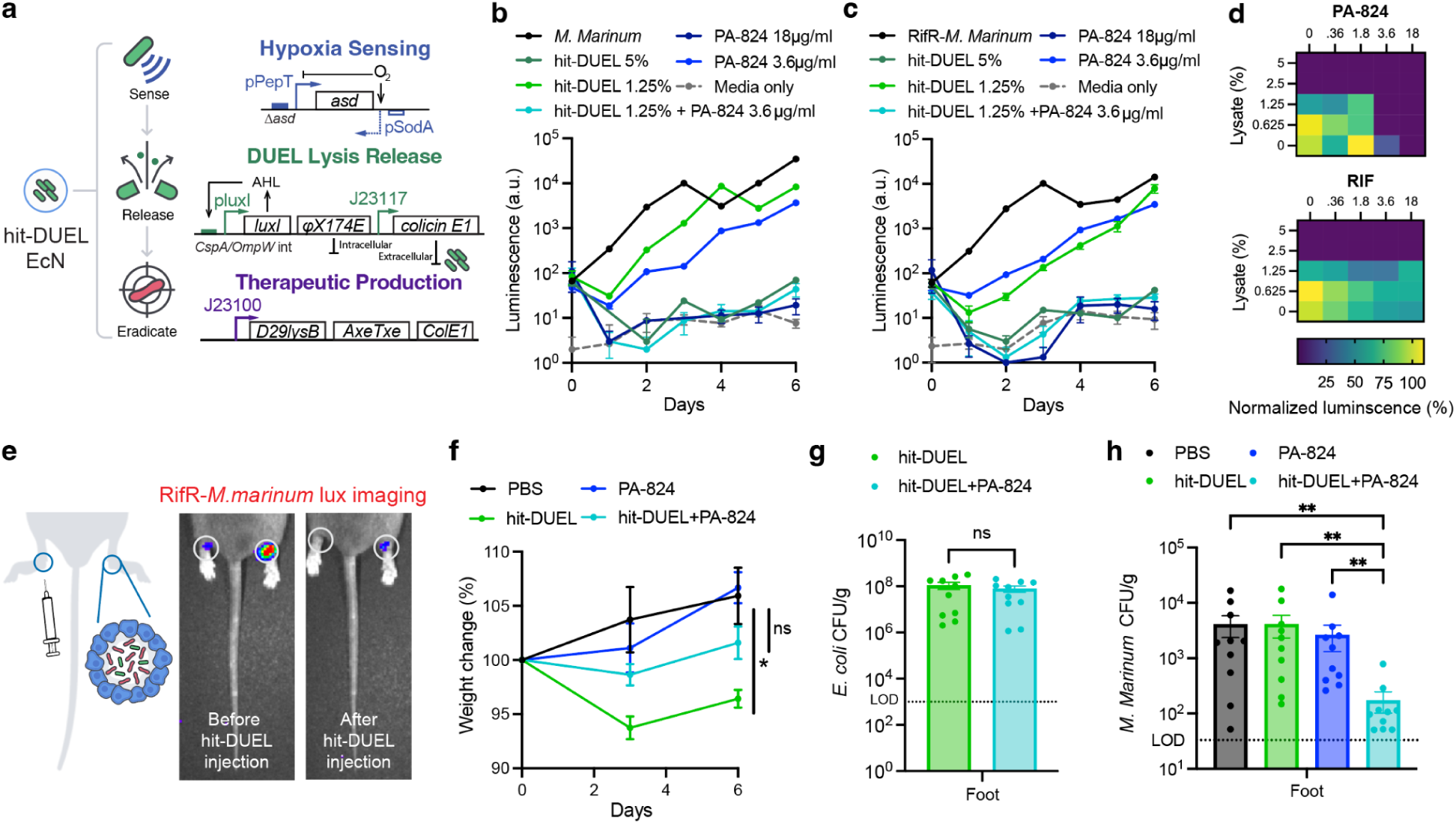
Engineered hit-DUEL enhances antibiotic activity against drug-sensitive and rifampicin-resistant mycobacteria. **a**, Schematic of hit-DUEL EcN: hypoxia-sensing circuit, integrated DUEL lysis, and therapeutic D29 LysB expression. **b-c,** Luminescence signal of *M. marinum* (*Mm*-lux) (**b**) and rifampicin resistant-*M. marinum* (RifR-*Mm*-lux) (**c**) in LB/7H9 1:1 medium over six days following treatment with hit-DUEL lysate, pretomanid (PA-824), or subtherapeutic combinations. High-dose of lysate (5%) or PA-824 (18 μg/mL) suppressed both drug-sensitive and RifR-*Mm*; however, the combination of subtherapeutic lysate (1.25%) and subtherapeutic PA-824 (3.6 μg/mL) also achieved sustained growth suppression through day 6. **d,** Dose-response heatmaps of day 6 RifR-*Mm*-lux luminescence across hit-DUEL lysate (0–5%) and different concentration of either PA-824 (top) or RIF (bottom). Lysate shifted the effective antibiotic dose required for growth suppression to lower concentrations, indicating synergistic potentiation. **e,** Schematic and representative IVIS image of RifR-*Mm*-lux-infected C3HeB/FeJ footpads before (day 0) and after (day 3) intra-lesional hit-DUEL injection. **f,** Body weight change following treatment with PBS, PA-824 (100 µM, ∼0.2 mg/kg), hit-DUEL, or hit-DUEL + PA-824 over 6 days. Combination treatment with PA-824 limited body weight loss relative to hit-DUEL alone over 6 days. **g,** Viable EcN recovered from footpad lesions at day 6 with or without PA-824 showed comparable bacterial persistence, indicating no antagonism between probiotic and antibiotic. **h,** Bacterial load of *M. marinum* CFU/g in footpad granulomas six days post-treatment. Combination therapy reduced burden compared to monotherapies or PBS control. Data represent mean ± s.e.m (b, c, f, g, h); n = 3 biological replicates (b, c, d), n = 5 mice (f), n = 10 feet (g, h). Statistical analysis was performed using two-way ANOVA (f), unpaired t-test (g), Kruskal-Wallis test (h); \**P* < 0.033, \*\**P <* 0.002, \*\*\**P <* 0.001; ns, not significant.

We next evaluated efficacy in a rifampicin-resistant (RifR) *M. marinum* (RifR-*Mm*-lux; *luxCDABE*) model. As in drug-susceptible *M. marinum*, neither bacterial lysate nor antibiotics alone achieved complete inhibition; however, combination treatment significantly suppressed bacterial growth (**Fig. 4c**). Notably, PA-824 combined with lysate eliminated RifR-*M. marinum*, and rifampicin (RIF) also achieved clearance when paired with low-dose bacterial lysates. A dose–response heatmap further demonstrated enhanced efficacy across conditions, with lysate lowering the effective antibiotic dose required for bacterial suppression (**Fig. 4d**). Co-culture of hit-DUEL with either *Mm*-lux or RifR-*Mm*-lux revealed that the combination of competition, therapeutic payload, and antibiotic treatment was important for sustaining robust mycobacterial growth suppression (**Supplementary Fig. 10**). These results demonstrate that hit-DUEL potentiates antibiotic activity and enables effective control of drug-resistant mycobacteria.

### hit-DUEL synergizes with antibiotics in vivo

To evaluate combinatorial efficacy in vivo, C3HeB/FeJ mice were infected with luminescent RifR-*M. marinum* (RifR-*Mm*-lux) through footpad injection. Mice with granulomas were then treated with hit-DUEL, with or without pretomanid (PA-824). Longitudinal IVIS imaging revealed reduced luminescence following hit-DUEL treatment, with further suppression observed under combination therapy (**Fig. 4e**). Treatment tolerability was assessed by body weight monitoring: hit-DUEL monotherapy induced a modest (∼5%) transient weight loss, whereas combination treatment with PA-824 (100 μM in 50 µL intra-lesional injection with hit-DUEL given twice, equivalent to ∼0.2 mg/kg based on injected volume and mouse body weight) maintained body weight comparable to PBS controls (**Fig. 4f**). Viable EcN were recovered at comparable levels from granulomas harvested 6 days after treatment with or without PA-824, indicating compatibility between probiotic and antibiotic (**Fig. 4g**). To confirm the preservation of lysis dynamics, we took bacterial colonies recovered after CFU plating and tracked their growth on a plate reader. We observed that all of the recovered colonies after plating on agar plates exhibited lysis oscillations indistinguishable from those of bacteria grown from the frozen glycerol stock, demonstrating maintenance of the integrated lysis operon and continued oscillatory behavior in vivo for at least 6 days after EcN injection (**Supplementary Fig. 11)**. Quantification of *M. marinum* burden indicated that combination therapy reduced *Mm* CFU by ∼20-fold relative to the untreated mice and outperformed both PA-824 and hit-DUEL monotherapy (**Fig. 4h**). Together, these results demonstrate that hit-DUEL suppresses drug-resistant mycobacterial infection and synergizes with pretomanid to achieve combination efficacy exceeding either monotherapy, supporting the translational potential of engineered probiotics as adjuncts to conventional TB treatment.

## Discussion

Engineered bacteria have demonstrated selective colonization of diseased physiological niches. Here, we extend this paradigm to infectious granulomas, exploiting their structural parallels with tumors, such as hypoxia, necrosis, and restricted vascularization, to enable probiotic colonization and localized antimycobacterial delivery. By integrating hypoxia-restricted growth, a dual-mode lysis circuit that suppresses evolutionary escape, and expression of a mycobacteriophage-derived enzyme, engineered EcN strains achieved suppression of drug-susceptible and rifampicin-resistant *M. marinum* in combination with subtherapeutic pretomanid, demonstrating proof-of-concept for a programmable, granuloma-targeted living therapeutic. This framework may generalize to the broader class of granulomatous infections (e.g. pathogens such as *Salmonella*, *Leishmania*, *Schistosoma*, *Histoplasma*) that similarly shelter pathogens within structured, hypoxic lesions.

A central question for engineered bacterial therapies is their translation to patients. Live bacterial therapeutics have established clinical precedent across multiple settings, most notably with *Mycobacterium bovis* BCG, which has been administered intravesically for bladder cancer since FDA approval in 1990. More recently, engineered strains of *Salmonella typhimurium* and *E. coli* Nissle 1917 (EcN) have demonstrated acceptable safety profiles in clinical studies, including rapid bloodstream clearance, lack of grade III/IV toxicities, and sustained localization within target tissues^19,20^. These approaches use attenuations to lipopolysaccharide structure for reduced immunogenicity, and auxotrophies for growth-restriction, enabling both intravenous and intratumoral delivery approaches. For example, early-generation *S. typhimurium* strain VNP20009 (*ΔmsbB*, *ΔpurI*) showed favorable safety in Phase I trials^40^, and next-generation strains such as Saltikva (IL-2–expressing *S. typhimurium*) have completed a Phase IIb study, where combination treatment with chemotherapy nearly doubled overall survival in pancreatic cancer patients^19^. These findings highlight the feasibility of engineered bacterial therapies and their capacity to function in combination with conventional treatments.

In this context, hypoxia-restricted EcN reduced off-target accumulation and improved the dose–toxicity relationship in vivo. This design provides a safety strategy that does not rely on permanent genetic deletions that can compromise bacterial fitness, while also functioning as an intrinsic containment mechanism, as loss of the target microenvironment limits bacterial persistence and promotes clearance. Additional layers, including alternative promoter–gene pairs or kill-switch circuits, could further enhance safety and enable repeat dosing for systemic administration. Our current studies are based on a cutaneous *M. marinum* model that recapitulates caseous necrosis but does not fully capture pulmonary *M. tuberculosis* granulomas. Extension to pulmonary infection models will further define performance in lung lesions and inform delivery strategies, such as targeting disseminated granulomas through inhalation approaches. Nonetheless, the modular architecture of this EcN platform supports multi-cargo expression and systematic payload screening of anti-mycobacterial or host-directed payloads, providing a foundation for iterative optimization toward a new therapeutic modality for tuberculosis and other granulomatous infections.

## Materials and Methods

### Bacterial strains and culturing

The main Mycobacterial strain used in this study was *M. marinum* (ATCC BAA-535). The host chassis was *E. coli* Nissle 1917 (EcN). The hypoxia-sensing derivative (hEcN) was constructed as previously described^36^. Full strain and plasmid information are listed in Supplementary Table 2.

*E. coli* strains were cultured in LB (Sigma-Aldrich) at 37°C, 225 r.p.m. (or on 1.5% LB agar at 37°C), with antibiotics selected per strain/plasmid (spectinomycin, 100 µg/mL; kanamycin, 50 µg/mL; chloramphenicol, 25 µg/mL; erythromycin, 50 µg/mL). Lysis-circuit strains were cultured in 0.4% glucose for <16 hours to suppress basal luxI expression. hEcN cultures were supplemented with 100 µg/mL diaminopimelic acid (DAP) to complement the Δasd knockout.

*Mycobacterium* strains were cultured in Middlebrook 7H9 broth (Millipore) with 10% ADC (Sigma-Aldrich), 4% glycerol and 0.05% Tween-80 (Sigma-Aldrich) to prevent clumping; kanamycin (50 µg/mL) was added for *M. marinum*-luxCDABE and hygromycin B (50 µg/mL) for *M. marinum*-tdTomato. Solid cultures were grown on 7H11 agar (Millipore) with 10% OADC (BD) and Myco PVNA antibiotic mix (Thermo Fisher) to suppress contamination, at 30°C, protected from light. *M. marinum* was grown 5–7 days to reach working density before any experiments.

*M. tuberculosis* Erdman (ATCC 35801) was cultured in complete 7H9-T medium composed of Middlebrook 7H9 base (BD Difco, Franklin Lakes, NJ) supplemented with 10% (v/v) oleic acid–albumin–dextrose–catalase (OADC). The OADC supplement consisted of oleic acid (0.5 g/L; Thermo Fisher Scientific, Waltham, MA), albumin (50 g/L; Fisher Scientific, Hampton, NH), dextrose (20 g/L; Fisher Scientific), and catalase (0.04 g/L; Thermo Fisher Scientific). The medium was further supplemented with sodium chloride (8 g/L; Fisher Scientific), glycerol (0.5% v/v; Fisher Scientific), and 0.05% Tyloxapol (Thermo Fisher Scientific). Kanamycin (50 μg/mL) was added during culture to maintain selection pressure, and the cultures were incubated at 37 °C with constant shaking at 150 rpm. Work with *M. tuberculosis* was conducted at BSL3 laboratory.

### In vitro 3D-granuloma spheroid platform

THP-1 monocytes were maintained in RPMI 1640 (Gibco) with 10% fetal bovine serum (FBS) and 1% penicillin/streptomycin at 37°C with 5% CO_2_. To differentiate macrophages, monocytes were treated with 100 ng/mL phorbol 12-myristate13-acetate (PMA; Sigma-Aldrich) for 24 hours, harvested by trypsinization (2–5 min) and mechanical scraping, then centrifuged (200 × *g*, 5 min) and resuspended in RPMI without PMA or antibiotics. Activated macrophages (1×10^6^ cells/well in 1 mL) were combined with *M. marinum*-tdTomato at an MOI of 1 or 0.1 in Aggrewell 400 24-well plates (STEMCELL Technologies) and centrifuged (100 × *g*, 3 min) to form spheroids at 30°C with 5% CO_2_. On day 2, spheroids were washed three times with 1× DPBS (70% media exchange per wash) and maintained in RPMI with 10 µg/mL gentamicin (Millipore Sigma). On day 3, EcN-sfGFP was added and co-incubated for 6 hours at 37°C, followed by three DPBS washes and resupplementation with gentamicin-containing RPMI.

### Microscopy

Spheroid images were acquired on an EVOS FL Auto 2 Imaging System (Celleste software). Time-lapse imaging of lysis-circuit dynamics used a Nikon Ti-E inverted microscope with an Okolab stage-top incubator (37°C, 5% CO_2_) and an Andor Zyla sCMOS camera, controlled by NIS-Elements software, with images acquired hourly. Phase-contrast images were taken at 10x magnification (50–200 ms exposure); GFP (sfGFP) and Texas Red (tdTomato) fluorescence were imaged at 10x, 70 ms exposure, 30% Lumencor Spectra-X intensity.

### DUEL lysis circuit construction

The DUEL plasmid was built on the synchronized lysing circuit (SLC) backbone with p15A origin of replication^41^. The colicin E1 (CE1) was synthesized (Eton Bioscience) based on the UniProt sequence P02978. Primers were designed to PCR the colicin E1 gene with different constitutive promoters with varying strengths from the Anderson promoter library BBa_J23119 (J23119, J23114 with 0.10 measured strength, and J23117 with 0.06 measured strength). The SLC backbone plasmid and colicin E1 insert were assembled (NEBuilder HiFi DNA Assembly) and transformed into a commercial electrocompetent 5-alpha *E.coli* comp cell (New England Biolabs). These constructs are referred to by the last three digits of the promoter designation for CE1: DUEL114 and DUEL117. The plasmid sequence was verified using whole-plasmid sequencing, and then transformed into working strains EcN or hEcN. The DUEL117 construct was chromosomally integrated into the EcN and hEcN strain genome at the *cspA* and *ompW* loci using the single-plasmid INTEGRATE system (pSPIN) with locus-specific crRNA guides ^34^; the resulting strain is referred to as i-DUEL. Correct integrants were confirmed by colony PCR and Sanger sequencing, and lysis behavior was checked with plate reader experiment.

### Therapeutic plasmid construction

Plasmids were constructed by Gibson assembly or restriction cloning. AMP-coding sequences were synthesized (Integrated DNA Technologies) based on published sequences (Supplementary Table 1) and cloned under the luxI promoter, replacing sfGFP in the ColE1 plasmid pTD103-sfGFP to generate the pTH05 inducible reporter plasmid^42^. The pTH05 backbone was linearized by PCR (primers pTH05_for/pTH05_rev), and each AMP gene was inserted by Gibson assembly and transformed into Mach1 cells (Invitrogen). For the most potent candidates, the luxI promoter was replaced with the constitutive J23100 promoter, and a C-terminal 6x-His tag with a GGSGG linker was added to the end of the coding DNA sequence. An Axe/Txe stabilization cassette was included for in vivo plasmid retention^43^.

The D29 LysB is a serine esterase with a Ser-Asp-His catalytic triad. The catalytic serine at position 82 was substituted with alanine (S82A)^32^ by PCR with a set of primers incorporating the intended point mutation, followed by Kinase-Ligase-DpnI (KLD) treatment (New England Biolabs), and transformation into *E. coli* 5-alpha cells (New England Biolabs). The complete open reading frame was sequence-verified by Whole-plasmid sequencing to confirm the intended mutation and exclude secondary substitutions.

### Lysate preparation for library screening

The pTH05 construct contains a pluxI promoter driving antimicrobial peptide (AMP) expression. Each EcN expressing AMP strains were grown overnight in LB with appropriate antibiotics, then subcultured 1:1000 into fresh LB and induced with N-acyl homoserine lactone (AHL, 100 nM) and grown overnight in a 37°C shaker. The cells corresponding to an optical density (OD_600_) of 1.0 in 1 mL were taken and centrifuged at 3,000 × *g* for 10 min, resuspended in 1 mL LB, and lysed by sonication (Fisherbrand Model 505 Sonic Dismembrator) using a 3 s on/off pulse cycle for a total duration of 1 min 30 s at 21% amplitude. Lysates were clarified by centrifugation at 3,000 × g for 10 min at 4°C, and supernatants were collected for immediate use in downstream screening assays.

### Western blot

Overnight cultures (5 mL) were grown and cells were harvested by centrifugation at 3,000 × *g* for 10 min and resuspended in 1 mL PBS. Cell density was normalized across samples to the same OD_600_ of 3 prior to a second centrifugation step. After a second centrifugation, pellets were lysed by freeze-thaw (liquid nitrogen, 5 min; 42°C water bath, 10 min) followed by chemical lysis using B-PER Bacterial Protein Extraction Reagent (ThermoFisher) with Halt Protease Inhibitor Cocktail. Soluble lysate was recovered by centrifugation, then mixed with 4× Bolt LDS sample buffer at a 1:3 ratio, and heated at 95°C for 5 min. Prepared samples can either be analyzed immediately or stored at −20°C for future use. The samples were resolved on Bolt 10% Bis-Tris Plus Mini Gels with 1× Bolt MES SDS Running Buffer (Thermo Fisher Scientific) at 200 V for 30 min. Gels were transferred to PVDF membranes using the iBlot 2 system (Thermo Fisher Scientific). Membranes were blocked in 5% non-fat dry milk in PBS-T (PBS, 0.05% Tween-20) for 1 hour at room temperature, then washed 2–3 times with PBS-T. Blots were incubated with anti-His primary mouse antibody (GenScript, A00186; 1:5000 dilution) for 30–60 min at room temperature. After three washes with PBST, membranes were incubated with anti-mouse HRP-conjugated secondary antibody (Cell Signaling, #7076, 1:1000 dilution) for 30–60 min, washed three times, and finally Western HRP substrate (Millipore, BLUR0100) was added right before imaging with ChemiDoc Imaging Systems (Bio-rad).

### Preparation of *M. marinum* for in vitro assay

*M. marinum* cultures were centrifuged at 3,000 × *g* for 10 min, resuspended in 5 mL fresh 7H9 media. To obtain the most single-cell in suspension, the cultures were sonicated in a water bath sonicator (Bransonic M1800) for 2–5 min. Following sonication, cultures were gently centrifuged at 150 × g for 1 min to pellet residual clumps, and the upper single-cell suspension was carefully collected. OD_600_ was adjusted to the desired number. The OD_600_ to CFU conversion was set to be OD_600_=1 is equal to 1×10^6^ CFU for downstream experiment.

### EcN lysate preparation and in vitro efficacy assays

Overnight cultures of EcN strains were grown from glycerol stocks with appropriate antibiotics and cultured for less than 16 hours. For lysate preparation, cultures were subcultured to OD_600_ = 0.01 in 5 mL LB without antibiotics (to avoid antibiotic carryover into downstream assays) and grown for 16 hours at 37°C with shaking. Cultures were centrifuged at 3,000 × *g* for 10 min at room temperature. Supernatants were carefully collected, centrifuged a second time, and then filtered through 0.22 μm sterile syringe PVDF filters (MilliporeSigma) into 14-mL conical tubes to remove residual cells. Supernatants were stored at 4°C to prevent protein degradation prior to use in downstream assays and can be used within 2 weeks.

For the plate setup, lysates were serially diluted two-fold (at 2x final concentration) across a 96-well plate (1.25–10% v/v). *M. marinum*-lux or RifR-*M. marinum*-lux (prepared as OD_600_ = 0.02 in 7H9) was added 1:1 to each dilution, yielding final concentrations of 0.625–5% lysate and OD_600_ = 0.01 mycobacteria. Reactions (100 µL, triplicate) were added to a white, clear-bottom 96-well plates (Corning), sealed with Breathe-Easy membrane (Sigma-Aldrich), and incubated at 30°C. OD_600_ and luminescence measurement were taken every 10 min for 20 hours with a plate reader (Infinite 200 PRO; Tecan), then once daily for 7 days.

### Antibiotic preparation for *in vitro* experiments

Pretomanid (PA-824; MedChemExpress) was dissolved in dimethyl sulfoxide (DMSO; Sigma-Aldrich) at 100 mM as a stock solution and stored at −20°C protected from light. Rifampicin (RIF; Sigma-Aldrich) was dissolved in DMSO at a stock concentration of 10 mM and stored at −20°C. Working concentrations were prepared by serial dilution in LB/7H9 (1:1) medium or deionized water immediately before each experiment, with a final DMSO concentration not exceeding 1% (v/v) in all assay wells. All antibiotic stocks were vortexed immediately before use, and working dilutions were prepared fresh before each experiment to minimize degradation.

### Combination treatment of lysate and antibiotic in vitro efficacy experiment

PA-824 or rifampicin (RIF) was serially diluted in DMSO/LB (1:100; final DMSO ≤1% v/v) to final concentrations of 1–50 µM (∼0.36–18 µg/mL). Lysate was independently serially diluted two-fold (final 1.25–10% v/v). Lysate and antibiotic dilutions were combined in a 96-well checkerboard format, then mixed 1:1 with *M. marinum*-lux or RifR-*M. marinum*-lux (OD_600_ = 0.02 in 7H9) to result in final concentrations of 0.625–5% lysate and 0.5–25 µM antibiotic at OD_600_ = 0.01 mycobacteria. Plate setup was prepared and read as described above; luminescence was normalized to the untreated mycobacterial control and reported as percent normalized luminescence.

### In vitro co-culture efficacy assay

Overnight cultures of each hypoxia-sensing strain (hEcN, hypoxic i-DUEL, hit-DUEL, and hit-DUEL MUT; see Supplementary Table 2) were centrifuged (3,000 × *g*, 10 min, room temperature) and resuspended in DAP-supplemented LB without antibiotics (OD_600_= 0.002). In a 24-well plate, each strain was tested against a two-fold dilution series of PA-824, with drug-free wells as controls. *M. marinum* or RifR-*M. marinum* (OD_600_ = 0.02) was added at equal volume (final OD_600_ = 0.01 mycobacteria, 0.001 EcN) in triplicate 100 µL wells (white, clear-bottom; Corning), sealed with Breathe-Easy membrane. Plates were incubated at 30°C; Mm-lux luminescence and EcN-GFP fluorescence were read every 10 min for 20 hours, then once daily for 7 days.

### M. tuberculosis in vitro assay

*M. tuberculosis* Erdman (L5 attB::Pleft-mScarlet) was cultured to an OD_600_ of 0.4–0.6, harvested by centrifugation (3,000 × *g*, 10 min), and resuspended in complete 7H9-T medium without kanamycin. EcN lysate or medium-only control was added to *M. tuberculosis* cultures at a final OD_600_ of 0.01. At the indicated time points (days 0, 1, 3, 5, 7, and 14), samples were diluted in PBS with 0.05% Tyloxapol and plated on 7H11 agar. Plates were incubated at 37°C with 5% CO_2_, and colony-forming units (CFU) were enumerated after a minimum of 21 days. All work with *M. tuberculosis* was conducted under BSL3 containment.

### Animal models

All animal experiments were approved by the Institutional Animal Care and Use Committee at Columbia University (protocols AC-AACF5755) and conducted in accordance with institutional guidelines. Animals were housed under standard conditions with a 12-hour light/dark cycle and ad libitum access to food and water. Mice were humanely euthanized upon loss of more than 20% initial body weight or upon recommendation of veterinary staff. Mice were randomized across treatment groups prior to experiments.

### *M. marinum* infection mouse model

Female C3HeB/FeJ or C57BL/6J mice (9–11 weeks; Jackson Laboratory) were used for granuloma induction. *M. marinum* was cultured for 7 days in 7H9 media (50 mL), washed three times in PBS (3,000 × *g*, 10 min), water-bath sonicated for 5 min to disperse clumps, and centrifuged gently (150 × *g*, 1 min) to collect the single-cell suspension. For intravenous infection, bacteria were adjusted to OD_600_ = 0.5, and 200 µL (1×10^5^ CFU) was injected into the tail vein using a restrainer, without anesthesia. For footpad infection, bacteria were adjusted to OD_600_ = 1, and 25 µL (2.5×10^4^ CFU) was injected per footpad. Granuloma formation was confirmed by IVIS (PerkinElmer) at days 7 and 14 and quantified in Living Image software; mice were randomized across groups to match baseline luminescence signals before treatment.

### EcN administration for in vivo experiments

EcN strains were grown overnight in LB with appropriate antibiotics/supplements (DAP, glucose), subcultured 1:100 without antibiotics, and grown to mid-log phase (OD_600_ = 0.1–0.4, targeting ∼0.2 to avoid premature lysis). Cultures were centrifuged (4°C, 3,000 × *g*, 10 min), washed three times in cold PBS, and resuspended to OD_600_ = 1 (WT EcN) or OD_600_ = 5 (hEcN). For intravenous injection, 200 µL was administered via tail vein using a restrainer. For retro-orbital injection, 50 µL at OD_600_ = 0.5 (WT EcN) was administered under isoflurane anesthesia. Mice were monitored by body weight and imaged by IVIS bioluminescence. For intra-lesional injection, 25 µL was injected directly into granuloma-bearing footpads under isoflurane anesthesia.

### Preparation of PA-824 for in vivo administration

For direct intra-lesion injection, PA-824 was diluted from DMSO stock into the prepared EcN suspension immediately before injection, at a final concentration of 100 μM ∼ 36 µg/mL (final DMSO ≤ 0.1% v/v).

### Biodistribution

At the study endpoint, mice were euthanized by CO_2_ asphyxiation. Tails (intravenous model) or feet (intra-lesional model) were surgically excised; tails were divided into four sections, avoiding direct excision through a granuloma. Liver and spleen were also collected for EcN safety profiling.

Whole organs (liver, spleen) were weighed and homogenized in 5 mL cold sterile PBS with 10% glycerol using a gentleMACS tissue dissociator (C Tubes, Miltenyi Biotec).

Granuloma tissue (tail sections, feet) was weighed and homogenized in sterile PBS with 10% glycerol (1 mL per tail section; 1.5 mL per foot) using a handheld tissue homogenizer (VWR 200; medium speed, 10 s). Samples should be kept on ice during processing.

For EcN enumeration, homogenates were serially diluted 10-fold in PBS and 5 µL was spotted on LB agar with the appropriate antibiotic/supplement; colonies were counted after overnight incubation at 37°C. For *M. marinum* enumeration, homogenates were diluted 25-fold and 100 µL was plated on 7H11 agar with 10% OADC and PVNA antibiotic, incubated at 30°C, and counted after 7–10 days. CFU/g was calculated from the dilution factor and tissue mass.

### Plate reader lysis oscillation assay for recovered colonies

Following biodistribution plating, individual EcN colonies recovered on LB agar were picked and inoculated into 1000 µL LB medium supplemented with DAP and 0.4% glucose and appropriate antibiotics in 14-mL round bottom tubes. The cultures were grown in a 37°C shaking incubator overnight along with the culture made from frozen glycerol stock. The cultures are then OD_600_ matched to 1 into 100µL LB with DAP and antibiotic but without glucose into black clear bottom 96-well plate. The plate was incubated at 37°C with continuous OD_600_ measurements taken every 10 minutes over 20 hours using a plate reader. Glycerol stock bacteria grown under the same conditions served as the positive control for lysis oscillation behavior.

### Histology and Immunohistochemistry (IHC)

Granuloma tissues were fixed in 10% neutral buffered formalin (VWR) for 24 hours, before transferring to 70% ethanol. All tissues were then sent to the Histology and Imaging Core at the University of Washington for paraffin embedding, sectioning, and staining with hematoxylin and eosin (H&E). For IHC, sections were additionally stained with anti-Lipopolysaccharide antibody (absolute antibody, WN1 222-5; 1:10,000 dilution with tris-EDTA + Tween 20 antigen retrieval) to detect EcN, and anti-Mycobacterium tuberculosis polyclonal antibody (Invitrogen, PA1-7231; 1:20,000 dilution with tris-EDTA + Tween20 antigen retrieval) to detect *M. marinum*, which shares cross-reactive antigenic epitopes with *E. coli*. Acid-fast staining was performed using the Ziehl-Neelsen method.

## Statistical analysis

Statistical tests were calculated in GraphPad Prism 10.0. Data are presented as mean ± s.e.m. unless otherwise noted. Specific tests, sample sizes, and significance thresholds are indicated in the respective figure legends. Normally distributed data were analyzed by unpaired t-test, one-way ANOVA with Tukey’s post-hoc multiple comparisons test, or two-way ANOVA with Tukey’s post-hoc multiple comparisons test. Non-normally distributed data were analyzed by Kruskal-Wallis test with Dunn’s multiple comparisons. Mice were randomized in different groups before experiments.

## Acknowledgments

We thank D. Tobin for providing the *Mycobacterium marinum* integrated luxCDABE and *Mycobacterium marinum* integrated tdTomato. We thank B. Fountaine and B. W. Johnson for their advice and histologic processing, immunohistochemistry, and imaging with the Histology and Imaging core at the University of Washington. We thank E. Ballister for his expertise and advice on working with mycobacterium species. **Funding:** This work was supported by NIH R21AI171819 (T.D.) and the Taiwan-Columbia Scholarship funded by the Ministry of Education of Taiwan and Columbia University (Y.-Y.C.). **Author Contributions:** T.H., Y.-Y.C. and T.D. conceived the study. T.H. and Y.-Y.C designed and constructed the system and designed in vitro experiments. Y.-Y.C., T.H., H.K., Y.J., and H.J. constructed the plasmid and strains used for in vitro and in vivo experiments. J.I. constructed the therapeutic library and assisted with time-lapse microscopy. Y.-Y.C. and T.H. processed mice samples for histology. T.H., Y.-Y.C., and Y.J. developed the 3D-granuloma spheroid platform. Y.J. and X.H. performed western blot for protein quantification. Y.-Y.C., H.K., Y.J., H.J., and M.R.R. performed the in vitro efficacy experiments. Y.-Y.C., T.H., C.C., H.K., Y.J., H.J., S.Z., M.R.R., and X.H. performed *M. marinum*-infected mouse in vivo experiments. G.G.M. and B.W. performed *M. tuberculosis* in vitro efficacy experiments. **Competing interests:** Y.-Y.C., T.H., and T.D. have filed a patent application with the U.S. Patent and Trademark Office related to this work. The other authors declare that they have no competing interests. **Data and materials availability:** All data associated with this study are available in the main text or the Supplementary Materials. **Additional information:** Supplementary Information is available for this paper. Correspondence and requests for materials should be addressed to Tal Danino.

