## Supplementary Information for "Engineered probiotics target granulomas to treat mycobacterial infection"

Supplementary Figures 1–11

Supplementary Tables 1–2

Supplementary Movie 1

### Supplementary Figures

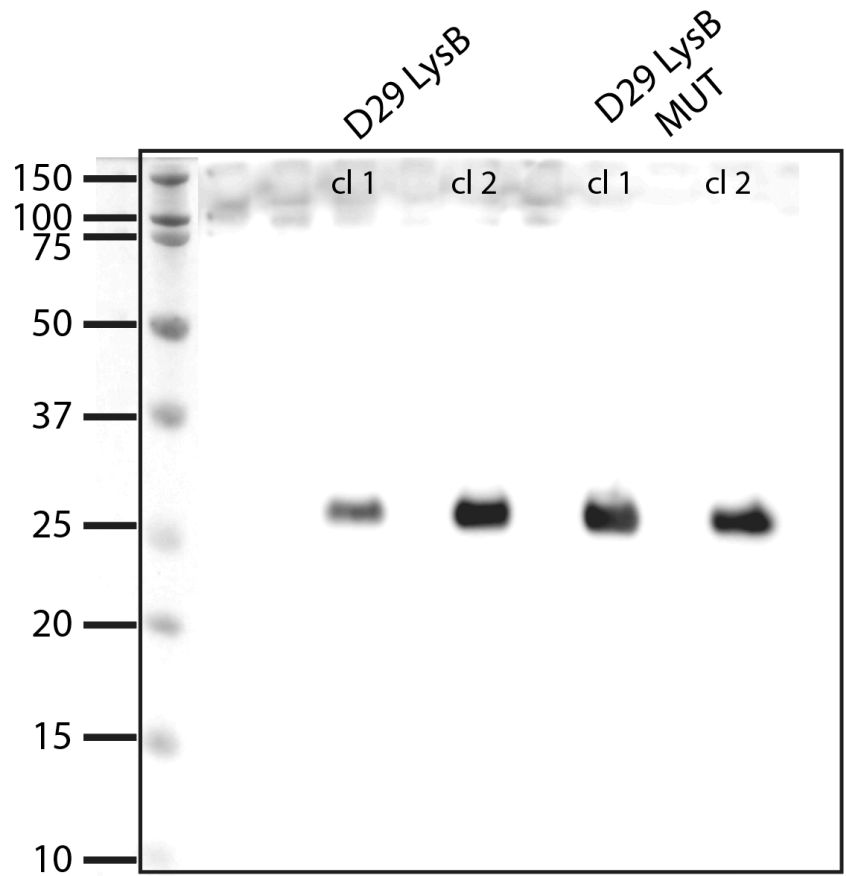

#### Supplementary Figure 1 | Expression of wild-type and catalytic mutant D29 LysB in EcN.

Western blot of EcN lysates probed with anti-Histag antibody. D29 LysB (wild-type) and the S82A catalytic mutant (D29 LysB MUT) were both detected at ~28.5 kDa, confirming equivalent expression. Molecular weight markers (kDa) indicated at left.

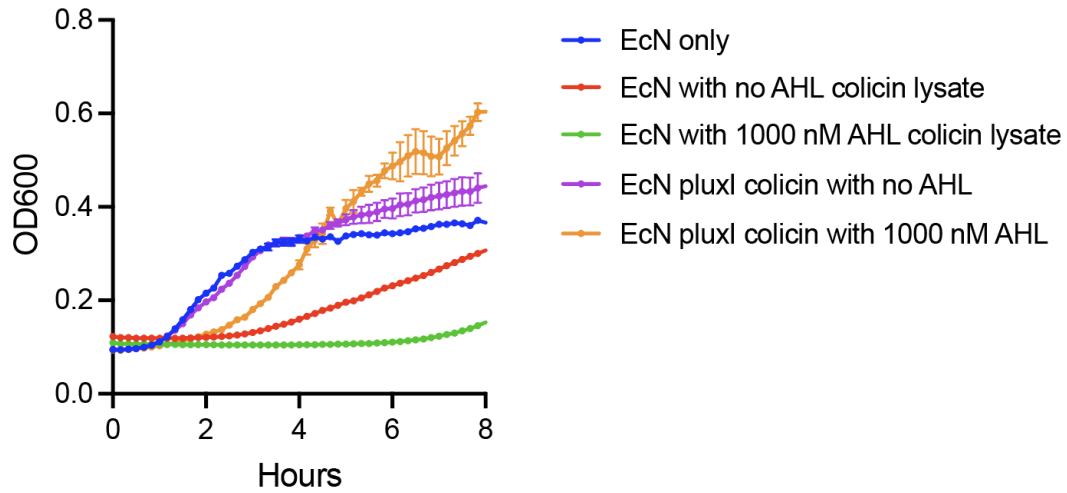

**Supplementary Figure 2 | Growth inhibition of EcN by extracellular Colicin E1 (CE1) but not intracellular expression.** Optical density at 600 nm ($OD_{600}$) growth curves of EcN exposed to CE1-containing lysate (extracellular delivery, quorum-induced; green) or expressing CE1 intracellularly under the *luxI* promoter, with (orange) or without (purple) AHL induction. EcN only control (dark blue) is shown for reference. Strong growth inhibition was observed only with extracellular CE1 delivery, indicating that CE1 must be released into the extracellular environment to exert its pore-forming activity. Data represent mean ± s.e.m.; $n = 3$ biological replicates.

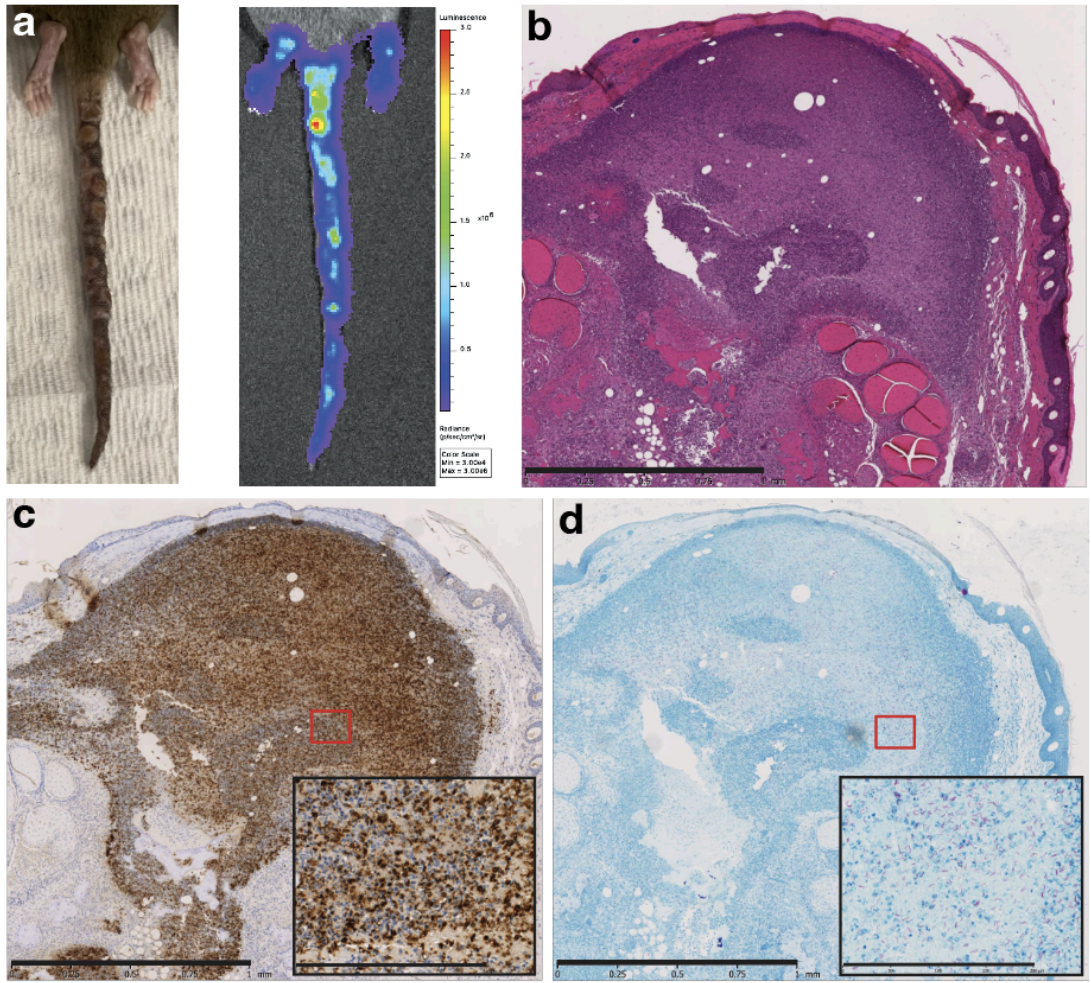

**Supplementary Figure 3 | Histological characterization of granulomas in intravenous *M. marinum*-infected mice.** **a**, Representative image and IVIS image of the infected tail from a C3HeB/FeJ mouse at 21 days post intravenous injection of *Mm* showing multiple nodular lesions and presence of live mycobacterium (FLuc, firefly luciferase). **b**, Hematoxylin and eosin (H&E) staining of tail tissue reveals large, organized granulomas with central caseous necrosis. **c**, Immunohistochemical staining with anti-*Mycobacterium tuberculosis* antibody showing dense mycobacterial bacilli within the necrotic core (brown). **d**, Acid-fast staining confirming *M. marinum* within granulomatous lesions (magenta). Scale bars as indicated. Scale bars (1 mm, inset: 200 μm) as indicated.

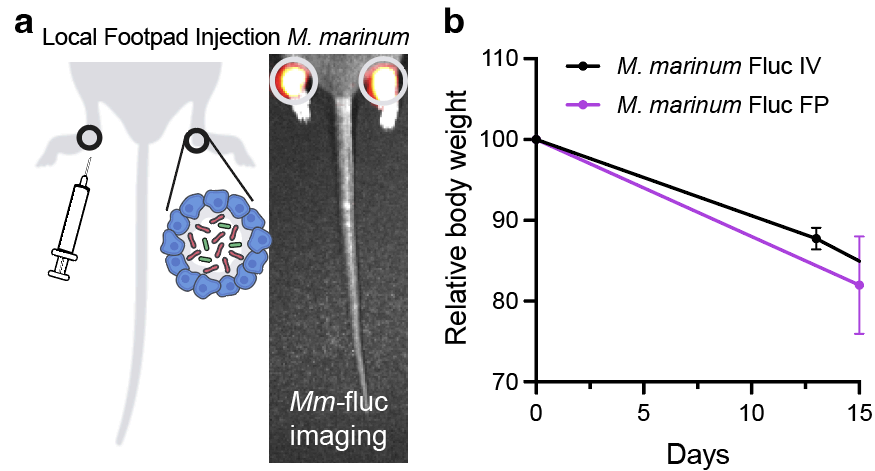

**Supplementary Figure 4 | Characterization of the direct footpad-injected *M. marinum***

**model. a,** Schematic and IVIS images of C3HeB/FeJ mice infected via footpad injection, showing *Mm*-FLuc (firefly luciferase) granuloma formation confined to the footpads after 14 days. **b,** Relative body weight comparison of mice receiving *Mm* via intravenous (IV) or footpad (FP) injection over 15 days. Data represent mean ± s.e.m.; n = 4 mice per group.

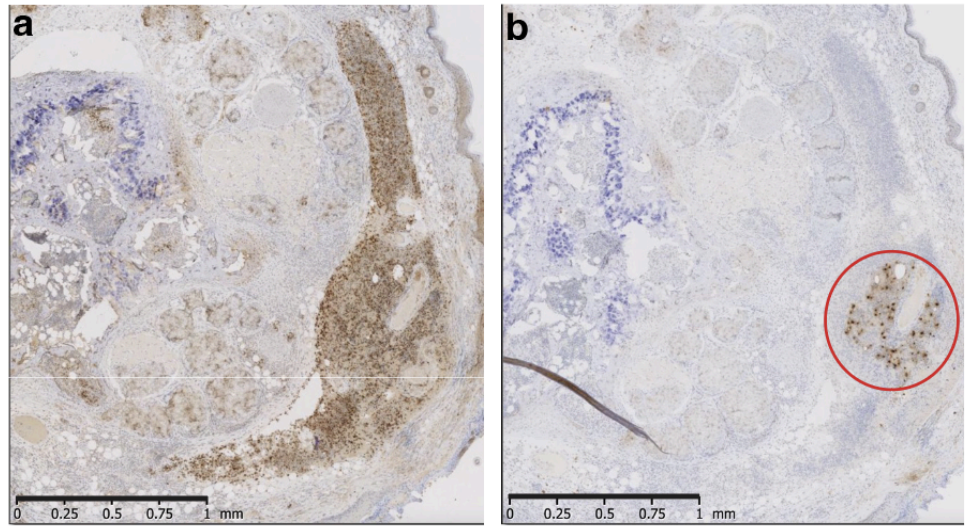

**Supplementary Figure 5 | Localization of systemically administered EcN within granulomas of *M. marinum*-infected mice.** Tissue sections from C3HeB/FeJ mice harvested 3 days after systemic EcN administration. **a**, Immunohistochemical staining with anti-mycobacterium polyclonal antibody showing mycobacterial signal within granulomatous lesions (brown signal). **b**, Anti-lipopolysaccharide (anti-LPS) staining detecting EcN in localized regions adjacent to and within granulomatous structures (red circle), demonstrating that systemically administered EcN traffics to and infiltrates mycobacterium-containing lesions. Scale bars (1 mm) as indicated.

#### *M. marinum* direct injection

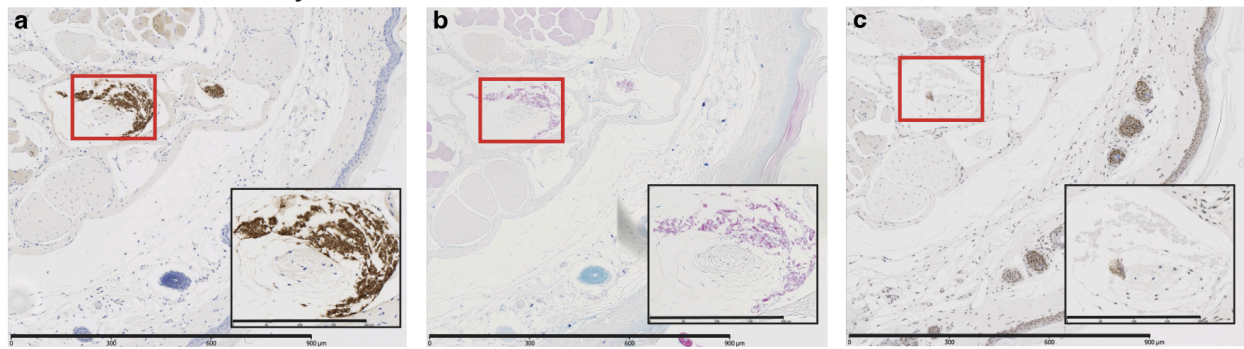

#### EcN direct injection

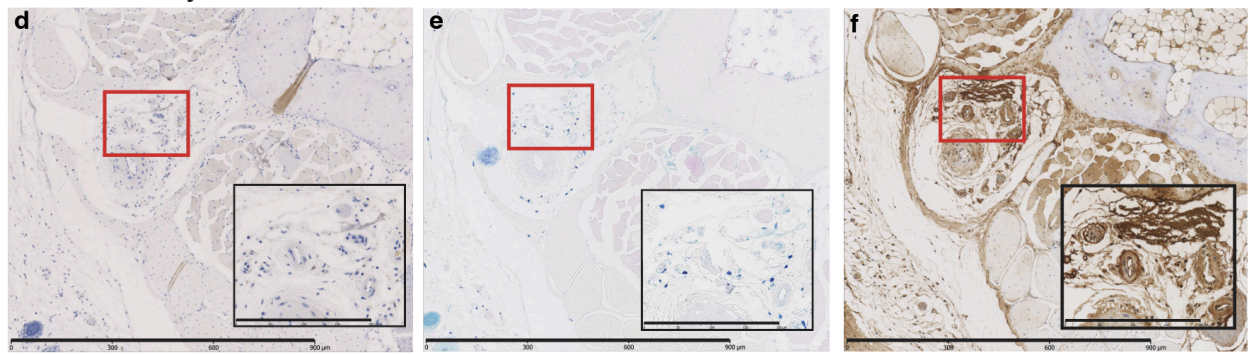

**Supplementary Figure 6 | Antibody specificity for mycobacterium and EcN detection *in vivo*.** Tail tissue injected with *M. marinum* alone (a–c) or EcN alone (d–f). **a**, Anti-mycobacterium immunohistochemistry labels mycobacterial aggregates (brown). **b**, Acid-fast staining confirms bacilli at the same sites (magenta). **c**, Anti-LPS staining shows no signal in Mm-injected tissue. **d,e**, Anti-mycobacterium and acid-fast staining show no signal in EcN-injected tissue. **f**, Anti-LPS selectively labels EcN (brown) in EcN-injected tissue. Insets show higher-magnification views of boxed regions; no cross-reactivity was detected between species-specific antibodies. Scale bars, 900 $\mu\text{m}$ (insets, 200 $\mu\text{m}$).

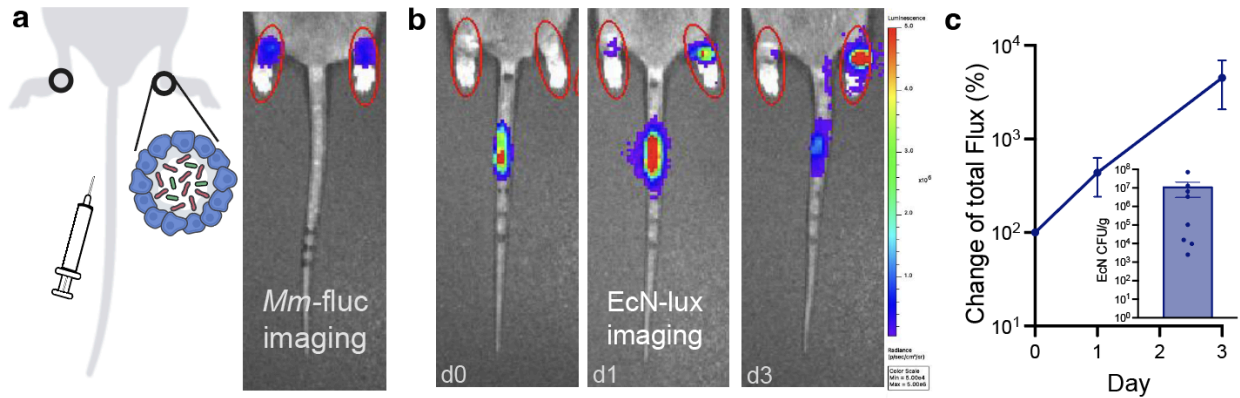

#### Supplementary Figure 7 | Systemically administered EcN localizes in footpad *M. marinum*-infected mice.

**a**, IVIS imaging showing *Mm*-FLuc (firefly luciferase) signals in infected footpads (circled). **b**, Representative EcN-lux (luxCDABE) images at days 0, 1, and 3 post-intravenous EcN injection, showing progressive accumulation of luminescent EcN signal specifically at *Mm*-infected footpads (circled) and not at uninfected sites. **c**, Quantification of EcN-lux luminescence signal in footpad over time, expressed as percent change in total flux relative to day 0. Inset: viable EcN CFU/g recovered from footpad at the endpoint day 3. Data are presented as mean ± s.e.m; n = 5 mice (c).

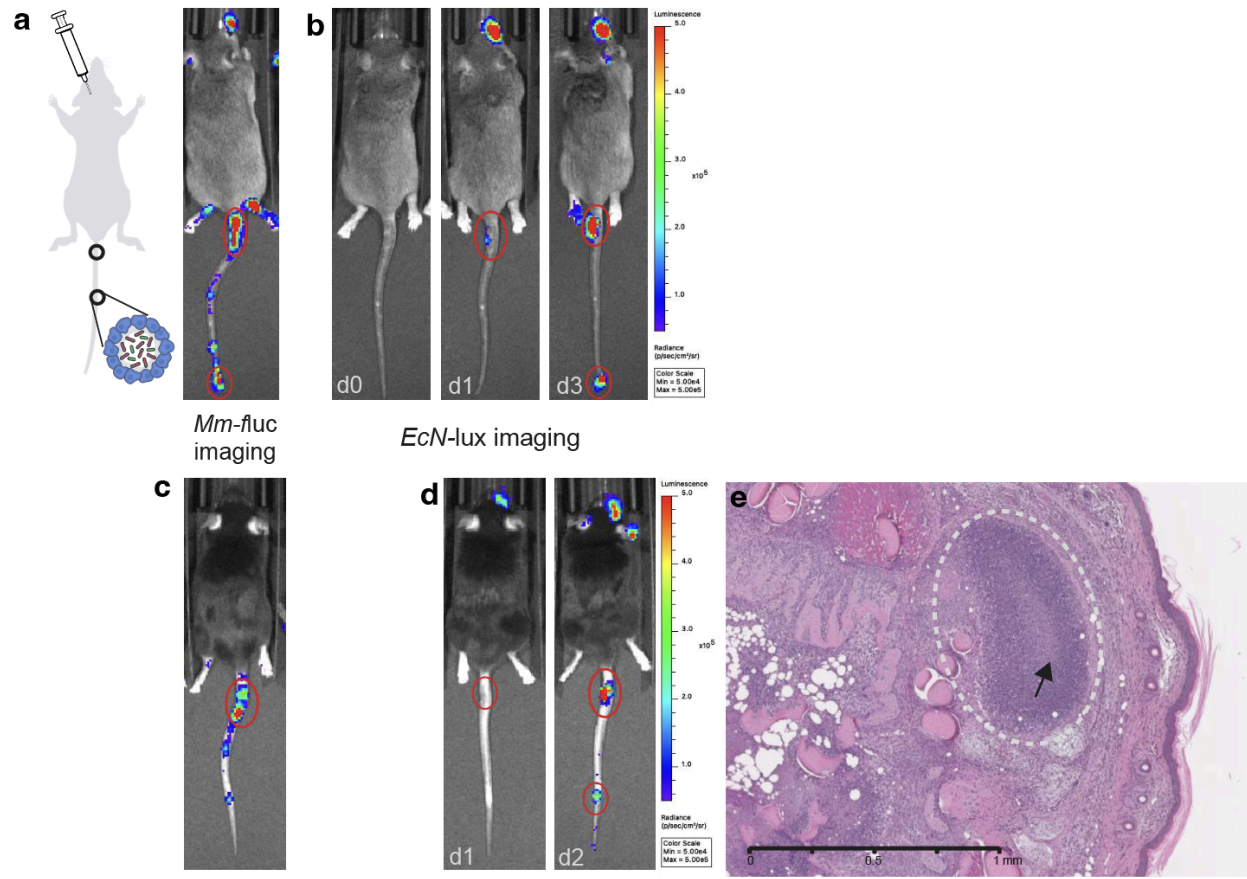

#### Supplementary Figure 8 | Granuloma colonization by EcN following retro-orbital delivery

**in C3HeB/FeJ and C57BL/6J mice. a,** IVIS image of a C3HeB/FeJ mouse at 21 days

post-intravenous injection of *Mm*-FLuc, showing disseminated bioluminescent lesions in the tail.

**b,** Representative EcN-lux (luxCDABE) images at days 0, 1, and 3 post-retro-orbital (RO) EcN injection, showing increasing luminescence signal localized to infected tail (circled), confirming granuloma colonization via this alternative delivery route. **c,** IVIS image of a C57BL/6J mouse at

21 days post-intravenous injection of *Mm*-FLuc, confirming granuloma formation in an

alternative host background. **d,** Representative EcN-lux images at days 1 and 2 post-RO

injection, showing EcN signal localized to the *Mm*-infected tail (circled). **e,** H&E staining of tail

tissue from a C57BL/6J mouse showing a large organized granuloma (circled) with central caseous necrosis (arrow), confirming the structural hallmarks of mycobacterial granulomas in this strain. Scale bar (1 mm) as indicated.

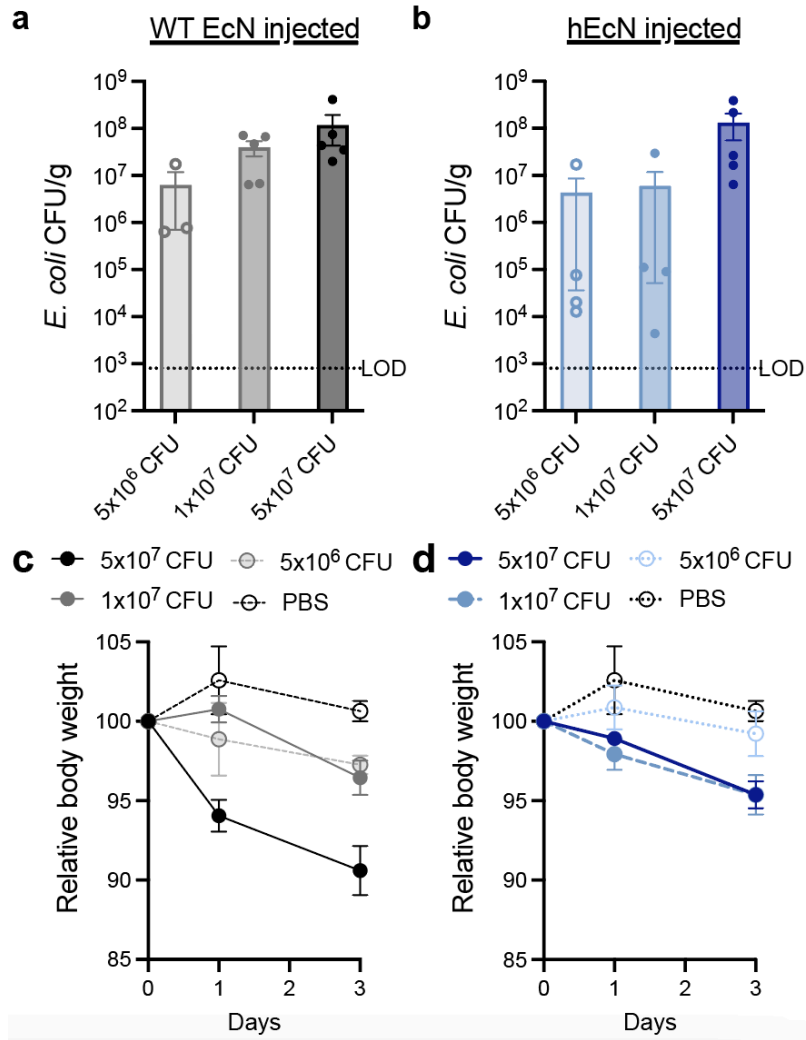

#### Supplementary Figure 9 | Dose-dependent colonization of granulomas and body weight

response following EcN administration in *M. marinum*-infected mice. Mice received $5 \times 10^6$,

$1 \times 10^7$, or $5 \times 10^7$ CFU of wild-type EcN or hypoxia-sensing hEcN intravenously. **a–b**,

Quantification of CFU/g recovered from tail granuloma from WT EcN (**a**) or hEcN (**b**), showing dose-dependent colonization. **c–d**, Relative body weight of mice following administration of WT

EcN (**c**) or hEcN (**d**) administration at the three doses over three days. Higher WT EcN doses

induced greater body weight loss; in contrast, hEcN at equivalent doses caused significantly less

weight loss while maintaining comparable granuloma colonization efficiency, demonstrating an

improved therapeutic index. Data are presented as mean ± s.e.m.; $n = 4$ mice per group.

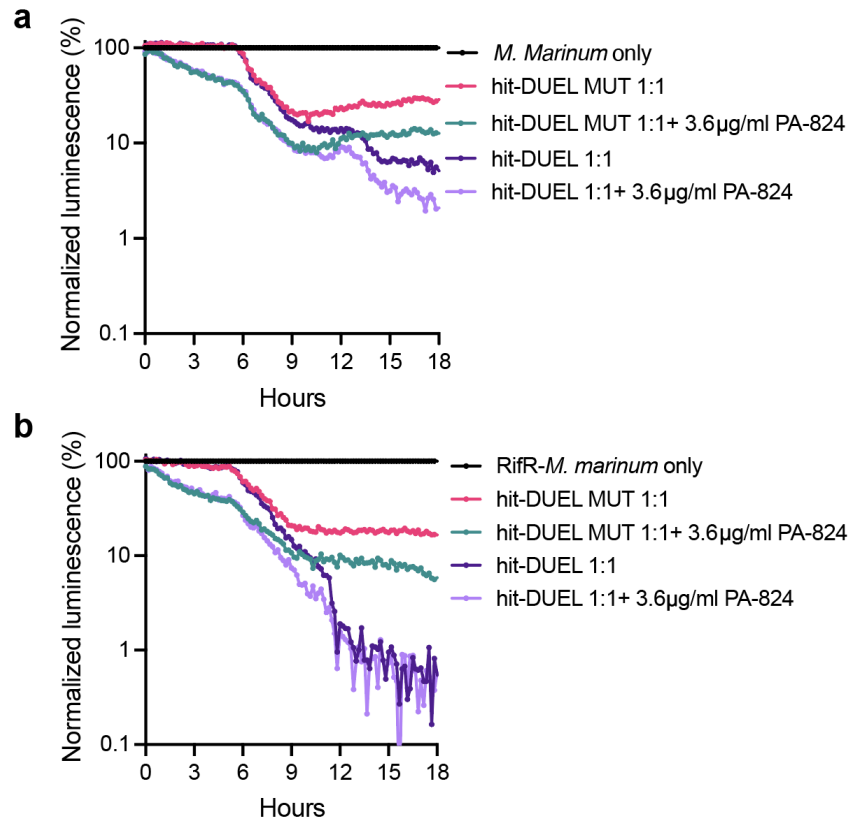

**Supplementary Figure 10 | Co-culture of hit-DUEL with *M. marinum* reveals payload- and competition-mediated suppression. a,** Normalized *Mm*-lux luminescence in 1:1 co-culture with hit-DUEL or hit-DUEL MUT, with or without subtherapeutic PA-824 (10 µM, ~3.6µg/mL). hit-DUEL MUT reduces luminescence relative to *M. marinum* alone through nutrient competition alone; hit-DUEL reduces it further via D29 LysB payload, and combination with PA-824 gives the strongest suppression. **b,** The same effect extended to Rif<sup>R</sup>-*Mm*-lux, showing that competition, therapeutic payload, and antibiotic combine to suppress rifampicin-resistant *M. marinum* (n = 3 biological replicates).

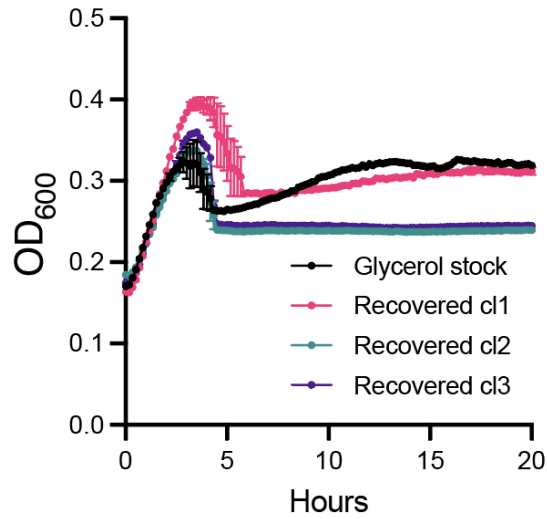

**Supplementary Figure 11 | hypoxia-sensing integrated i-DUEL lysis behavior after mouse experiment.** OD<sub>600</sub> growth curves of EcN colonies recovered from plating of footpad granulomas at day 6 post-treatment (Recovered cl1–cl3; magenta, teal, and purple) compared to bacteria grown directly from frozen glycerol stock (black). All cultures were OD<sub>600</sub> matched with a starting OD<sub>600</sub> = 1. All recovered colonies exhibit characteristic lysis oscillations, confirming maintenance of the integrated i-DUEL lysis operon and circuit stability following one week of in vivo colonization. Data represent mean ± s.e.m.; n = 3 biological replicates. Each colony is from a different mouse.

### Tables

**Supplementary Table 1.** Candidates for AMP

| Name | Target | Category | Mechanism of action | Origin | Citation |
| --- | --- | --- | --- | --- | --- |
| Bacteriocin E50-52 (BCN5) | plasma membrane | Bacteriocin | Pore-forming | *Enterococcus faecium* | 44 |
| BTCU-lysB | mycolic acid + peptidoglycan | Endolysin | Esterase (cleaves the ester bond between mycolic acid and arabinogalactan-peptidoglycan), lipolytic | BTCU (Phage) | 45 |
| Buforin IIb | DNA | Cationic AMP | DNA/RNA binding by trespassing the cell non-lytic | *Bufo gargarizans* (Frog) histone H2A | 46 |
| Cecropin X | plasma membrane | Cationic AMP | pore-forming | *Hyalophora cecropia* (Insect) | 46,47 |
| D29 LysB | mycolic acid + peptidoglycan | Endolysin | Esterase (cleaves the ester bond between mycolic acid and arabinogalactan-peptidoglycan), lipolytic | D29 (Phage) | 32,48 |
| HNP-1 | plasma membrane, DNA | Defensins (cationic peptides) | Pore-forming, macromolecule synthesis (secondary), chemotactic, modulate cytokine production | Human (Mammal), neutrophil granules | 49 |
| Magainin | plasma membrane | Cationic AMP | Pore-forming | *Xenopus laevis* (Frog) | 46,47 |
| Mastoparan | plasma membrane | Cationic AMP | Pore-forming | *Pseudopolybia vespiceps testacea* (wasp venom) | 46,47,50 |
| PIN 2 [14] | plasma membrane | Cationic AMP | Pore-forming | *Pandinus imperator* (scorpion) | 51 |
| Pleurocidin | plasma membrane | AMP | Pore-forming, DNA/RNA/protein synthesis interruption | *Pseudopleuronectes americanus* (Fish) | <sup>52</sup> |
| PR-39 | DNA/Protein | PR-AMP | Macromolecular synthesis(DNA/Protein synthesis interrupt) | Porcine (Mammal) | <sup>53</sup> |
| SEQ1 | plasma membrane | Cationic AMP | Pore-forming | Synthetic | <sup>54</sup> |
| SEQ11 | plasma membrane | Cationic AMP | Pore-forming | Synthetic | <sup>54</sup> |
| SEQ43 | plasma membrane | Cationic AMP | Pore-forming | Synthetic | <sup>54</sup> |
| SEQ5 | plasma membrane | Cationic AMP | Pore-forming | Synthetic | <sup>54</sup> |
| TDMH | mycolic acid | Serine esterase | Esterase (cleaves the ester bond of trehalose dimycolate) inducing cell lysis | *M. smegmatis* (Bacteria) | <sup>55,56</sup> |

**Supplementary Table 2.** Main bacterial strains and plasmids used in this study.

| Identifier | Bacterial Strains | Plasmids (ORI/promoter) | Relevant Features | Figure |
| --- | --- | --- | --- | --- |
| EcN | *E. coli* Nissle 1917 | N/A | N/A | 1 |
| EcN-GFP | *E. coli* Nissle 1917 | sc101/J23118 | Reporter GFP gene expressed under J23118 promoter | 1, 2 |
| *M. marinum*-luxCDABE (*Mm*-lux) | *Mycobacterium marinum* | integrated | Reporter luxCDABE gene expressed | 1 |
| *M. marinum*-tdTomato (*Mm*-tdTomato) | *Mycobacterium marinum* | integrated | Reporter tdTomato gene expressed | 1 |
| D29 LysB | *E. coli* Nissle 1917 | ColE1/J23100 | Therapeutic plasmid producing D29 LysB | 1 |
| D29 LysB MUT | *E. coli* Nissle 1917 | ColE1/J23100 | Therapeutic plasmid control producing mutated D29 LysB S82A | 1 |
| *M. tuberculosis* (*M. tb*) | *Mycobacterium tuberculosis* Erdman | L5 attB::Pleft* mScarlet (addgene #169410) | Virulent *Mtb* with RFP reporter plasmid | 1 |
| SLC | *E. coli* Nissle 1917 | p15a/pluxI<br>ColE1/J23119 | Synchronized lysis circuit with quorum-sensing promoter expressing ϕX174 E<br>Reporter GFP gene expressed under J23119 promoter | 2 |
| DUEL114 | *E. coli* Nissle 1917 | p15a/pluxI and J23114<br>ColE1/J23119 | DUEL lysis circuit with quorum-sensing promoter expressing ϕX174 E and constitutive J23114 expressing colicin E1<br>Reporter GFP gene expressed under J23119 promoter | 2 |
| DUEL117 | *E. coli* Nissle | p15a/ | DUEL lysis circuit with | 2 |
|  | 1917 | pluxI and J23117<br>ColE1/J23119 | quorum-sensing promoter expressing ϕX174 E and constitutive J23117 expressing colicin E1<br>Reporter GFP gene expressed under J23119 promoter |  |
| Integrated DUEL (i-DUEL) | *E. coli* Nissle 1917 | Integrated at CspA and OmpW/pluxI and J23117<br>ColE1/J23119 | DUEL117 construct integrated into EcN<br>Reporter GFP gene expressed under J23119 promoter | 2 |
| *M. marinum*-Firefly Luciferase (*Mm*-FLuc) | *Mycobacterium marinum* | integrated | Reporter firefly luciferase gene expressed to track *M. marinum* in vivo | 3 |
| EcN luxCDABE (EcN-lux) | *E. coli* Nissle 1917 | pUC/phelp | Reporter luxCDABE gene expressed to track EcN in vivo | 3 |
| Hypoxia EcN (hEcN) | *E. coli* Nissle 1917Δ*asd* | integrated/pPepT | *asd* gene expressed under pPepT promoter | 3 |
| Hypoxia EcN luxCDABE (hEcN-lux) | *E. coli* Nissle 1917Δ*asd* | integrated/pPepT<br>ColE1/phelp | *asd* gene expressed under pPepT promoter<br>Reporter luxCDABE gene expressed to track hEcN in vivo | 3 |
| Hypoxia i-DUEL | *E. coli* Nissle 1917Δ*asd* | integrated/pPepT<br>Integrated at CspA and OmpW/pluxI | *asd* gene expressed under pPepT promoter<br>DUEL117 construct integrated into hypoxia EcN | S11 |
| hit-DUEL | *E. coli* Nissle 1917Δ*asd* | integrated/pPepT<br>Integrated at CspA and OmpW/pluxI | *asd* gene expressed under pPepT promoter<br>DUEL117 construct integrated into hypoxia EcN | 4 |
|  |  | ColE1/J23100 | Therapeutic plasmid producing D29 LysB |  |
| hit-DUEL MUT | *E. coli* Nissle 1917Δ*asd* | integrated/pPepT<br>Integrated at CspA and OmpW/pluxI and J23117<br>ColE1/J23100 | *asd* gene expressed under pPepT promoter<br>DUEL117 construct integrated into hypoxia EcN<br>Therapeutic plasmid producing mutated D29 LysB S82A | 4 |
| Rifampicin Resistant-*M. Marinum*-luxCDABE (RifR-*Mm*-lux) | *Mycobacterium Marinum* | integrated | Reporter luxCDABE gene expressed and serially cultured in rifampicin antibiotic to select for rifampicin-resistant colony. | 4 |

### **Movie**

**Supplementary Movie 1.** Timelapse video of DUEL117. EcN with plasmid DUEL117 lysis circuit and GFP reporter payload on an agar pad. Cell death caused by lysis will turn red due to propidium iodide (PI) staining, and then the GFP (green) payload will be released.

nissle GFP+117 c1(series 12).mov
